# IGF1 Receptor Regulates Upward Firing Rate Homeostasis via the Mitochondrial Calcium Uniporter

**DOI:** 10.1101/2021.12.11.469196

**Authors:** Maxim Katsenelson, Ilana Shapira, Eman Abbas, Boaz Styr, Saba Aïd, Martin Holzenberger, Silvio Rizzoli, Inna Slutsky

## Abstract

Regulation of firing rate homeostasis constitutes a fundamental property of central neural circuits. While intracellular Ca^2+^ has long been hypothesized to be a feedback control signal, the molecular machinery enabling network-wide homeostatic response remains largely unknown. Here we show that deletion of insulin-like growth factor-1 receptor (IGF1R), a well-known regulator of neurodevelopment and ageing, limits firing rate homeostasis in response to inactivity, without altering the baseline firing rate distribution. Disruption of both synaptic and intrinsic homeostatic plasticity contributed to deficient firing rate homeostatic response. At the cellular level, a fraction of IGF1Rs was localized in mitochondria with the mitochondrial calcium uniporter complex (MCUc). IGF1R deletion suppressed spike burst-evoked mitochondrial Ca^2+^ (mitoCa^2+^) by weakening mitochondria-to-cytosol Ca^2+^ coupling. MCUc overexpression in IGF1R-deficient neurons rescued the deficits in spike-to-mitoCa^2+^ coupling and firing rate homeostasis. Our findings highlight IGF1R as a key regulator of the integrated homeostatic response by tuning mitochondrial temporal filtering. Decline in mitochondrial reliability for burst transfer may drive dysregulation of firing rate homeostasis in ageing and brain disorders associated with aberrant IGF1R / MCUc signaling.

## Introduction

Neural circuits are composed of a large number of dynamic elements at various levels of organization. The operation of a neuronal circuit depends on the interaction between the intrinsic properties of the individual neurons and the synaptic interactions that connect them into functional ensembles. While some aspects of synaptic and spiking activity are dynamic, others show remarkable stability over long time periods (1). Despite a large variability in synaptic and intrinsic parameters, firing rate distributions and their mean firing rate (MFR) are maintained at a specific set-point value during ongoing spontaneous activity. MFRs are typically restored even in the presence of large perturbations to activity rates and patterns. The renormalization of MFRs to a set-point value has been observed in response to activity-dependent perturbations in cortical and hippocampal networks grown *ex vivo* (2–6) and to sensory deprivation in V1 cortex *in vivo* (7–10). MFR homeostasis can achieved by a wide repertoire of homeostatic processes, including adjustments at the level of synaptic strength, intrinsic excitability, and excitation-to-inhibition balance (11–13). Dysregulation of homeostatic plasticity has been proposed to drive synaptic and cognitive deficits in distinct brain disorders, including neurodevelopmental disorders (14, 15) and neurodegenerative disorders like Alzheimer’s disease (16, 17).

Despite significant progress in our understanding of neuronal homeostasis, how firing rates are sensed and how this information is converted into feedback signaling remain largely unknown. Intracellular somatic Ca^2+^ has been proposed to serve as a proxy of spiking activity because of their tight coupling and is modeled as a feedback control signal (18, 19). According to these models, deviations from a specific target Ca^2+^ value induce changes in effector proteins that result in renormalization of firing properties to a set point value. Indeed, somatic cytosolic Ca^2+^ (cytoCa^2+^) in excitatory neurons returns to a set-point value following sensory deprivation *in vivo* (9, 20) and following neuronal inactivity *ex vivo* (4). However, the molecular mechanisms that maintain cytoCa^2+^ around a target value and mediate the coordinated homeostatic response to stabilize firing rate distributions are unknown.

The insulin-like growth factor-1 receptor (IGF1R) signaling is a well-known, evolutionary-conserved pathway regulating brain development (21, 22) and ageing (23–26). In the central nervous system, IGF1/IGF1R signaling is critical for experience-dependent synaptic and neuronal plasticity in sensory cortices (27–29), adult neurogenesis (30–32), synaptic vesicle release (33–35) and neuronal excitability (35, 36). Moreover, IGF1R is an important modulator of mitochondrial Ca^2+^ (mitoCa^2+^) levels in hippocampal neurons (34). As mitochondria are involved in neuronal Ca^2+^ homeostasis (37), we have hypothesized that IGF1R is necessary for the integrated homeostatic response to activity perturbations.

To test this hypothesis, we examined here how IGF1R affects spike-to-Ca^2+^ transfer functions at mitochondrial and cytosolic compartments in soma of excitatory hippocampal neurons and how it influences homeostatic MFR response to inactivity at the network level. Our results demonstrate that somatic mitochondria selectively uptake cytoCa^2+^ evoked by spike bursts, but not by single spikes. Deletion of IGF1R weakened spike-to-mitoCa^2+^ coupling by decreasing mitoCa^2+^-to-cytoCa^2+^ coupling efficiency during periods of spike bursts. A decrease of burst-evoked mitoCa^2+^ transients in IGF1R-KO neurons was associated with transcriptional down-regulation of several members of the mitochondrial calcium uniporter (MCU) complex (MCUc). As a result, MFR compensation was impaired in response to inactivity in IGF1R-deficient neurons. Overexpression of MCUc in IGF1R-KO neurons rescued both mitoCa^2+^ and MFR homeostatic response. These results provide direct evidence for the critical role of IGF1R / MCUc signaling in upward MFR homeostasis at the population level in hippocampal networks.

## Results

### IGF1R deficiency limits homeostatic compensation of mean firing rate to inactivity

To explore the role of mitoCa^2+^ in MFR homeostasis, we first tested whether IGF1R is necessary for the homeostatic regulation of firing rate distributions and their means. For this, we used multi-electrode arrays (MEAs) for long-term recordings of spiking activity at the same cultured hippocampal neurons during baseline and throughout two days of activity perturbation. The GABA_B_ receptor agonist baclofen (Bac) was used as a persistent inhibitory perturbation (4). For conditional deletion of IGF1Rs, we used IGF1R^fl/fl^ mice (38) with a viral delivery of Cre-recombinase. We used adeno-associated virus (AAV) under the general promoter CBAP (AAV1/2-CBAP-Cre-Cerulean), thus creating IGF1R knockout (IGF1R-KO) networks (Fig. S1). For control (Ctrl) experiments, IGF1R^fl/fl^ cultures were infected with AAV1/2-CBAP-Cerulean. As expected from our early work (4, 6), Ctrl networks displayed a pronounced suppression of activity by baclofen (10 µM) that was restored to network’s baseline MFR after a period of two days (Fig. 1A–C). Conversely, the average firing rates of IGF1R-KO networks exhibited only partial (~45%) recovery subsequent to baclofen application (Fig. 1D–F). Moreover, distributions of Ctrl single-unit firing rates were indistinguishable before and two days after baclofen application (Fig. 2A). While baclofen effects were variable per single unit in Ctrl conditions (Fig. 2B), on average per-unit change was not significant following baclofen application (Fig. 2C). In contrast, MFR distribution was left-shifted following baclofen application in IGF1R-KO neurons (Fig. 2D) due to incomplete single-unit recovery (Fig. 2E,F). These experiments point towards IGF1R as a necessary component of homeostatic MFR recovery from inactivity. Notably, IGF1R deletion did not affect baseline firing rates (Fig. 2G) and patterns (Fig. 2H,I and Fig. S2).

**Figure 1.**
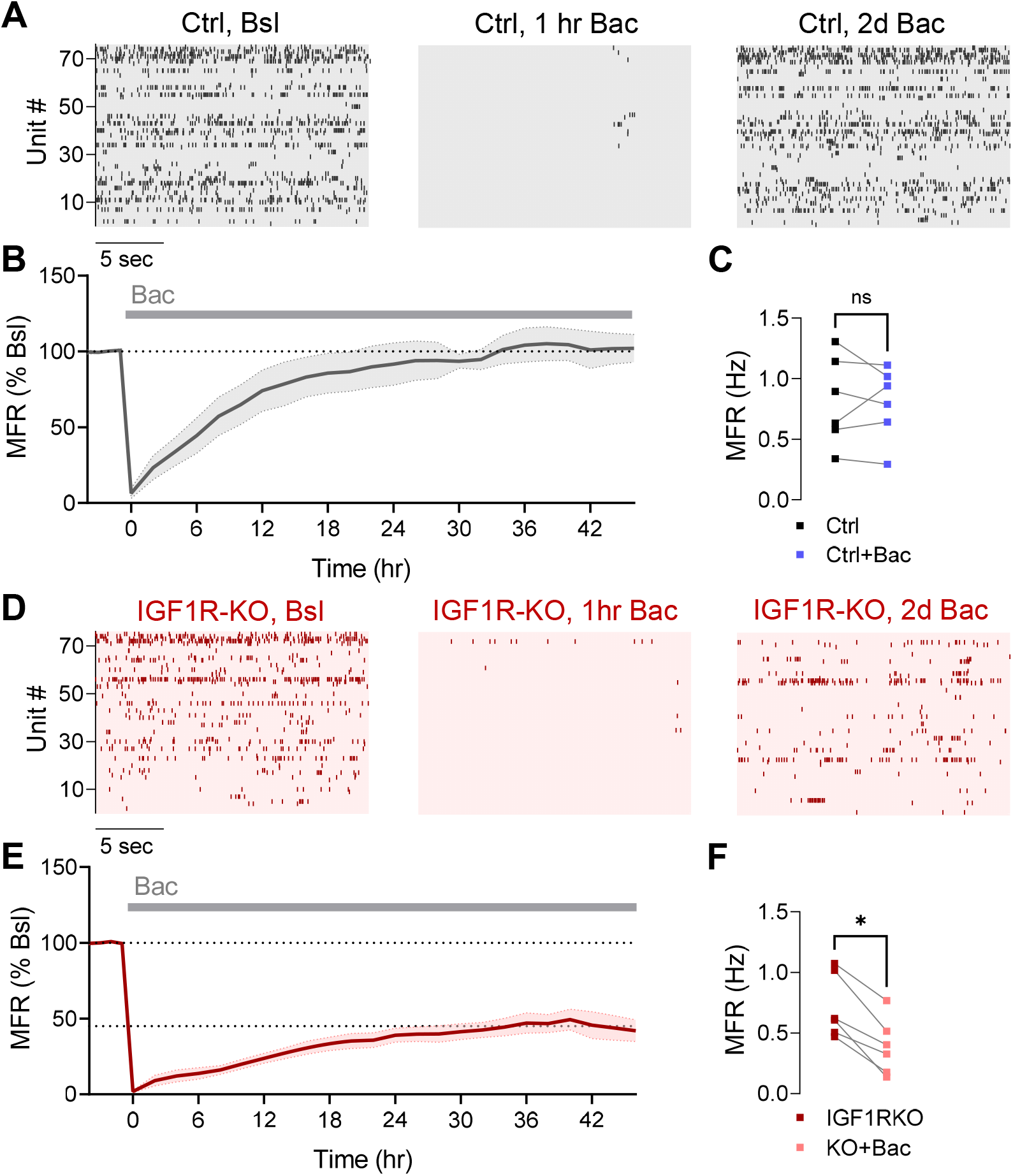
IGF1R deletion limits mean firing rate homeostasis at the network level. (*A*) Raster plots of the same neurons during baseline (Bsl, *Left*), acute application of 10 µM baclofen (Bac, *Center*), and following two days of Bac (*Right*) in Ctrl (Ctrl) cultures. (*B*) A typical MFR homeostatic response to chronic inactivity induced by 10 µM baclofen in Ctrl cultures, showing renormalization to the baseline network MFR (6 experiments, 535 units). (*C*) Summary of full MFR recovery following two days of Bac (Bac_2d_) in Ctrl networks (*P* = 0.68, each point represents individual experiment, same data as *B*). (*D*) Raster plots of the same neurons during baseline (*Left*), acute application of Bac (*Center*), and following Bac_2d_ (*Right*) in IGF1R-KO culture. (*E*) MFR homeostatic response was limited by IGF1R-KO (6 experiments, 371 units). (*F*) Summary of partial recovery of MFRs following Bac_2d_ in IGF1R-KO networks (*P* = 0.0313, each point represents individual experiment, same data as *E*). Wilcoxon test (*C,F*). Error bars indicate SEM. ns, not significant, \**P* < 0.05.

**Figure 2.**
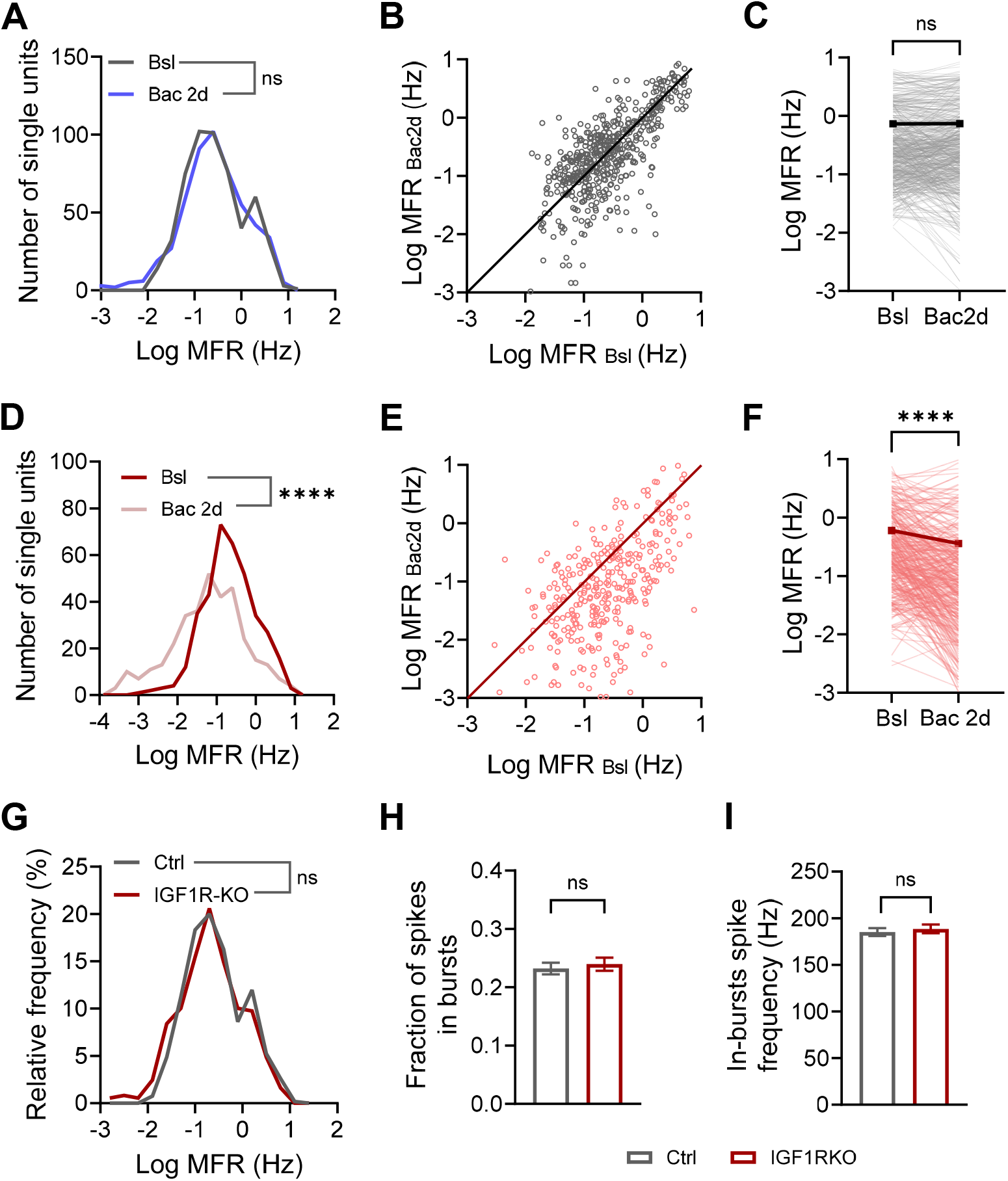
Deletion of IGF1R limits firing rate homeostasis, without affecting basal firing rate and pattern. (*A-C*) Effect of Bac_2d_ on 535 single units in Ctrl cultures. (*A*) Firing rate distributions were unchanged following Bac_2d_ (*P* = 0.23). (*B*) Changes in MFR per neuron. Line indicates no change. (*C*) MFRs were unchanged following Bac_2d_ (*P* = 0.79). (*D-F*) Effect of Bac_2d_ on 371 single units in IGF1R-KO cultures. (*D*) Firing rate distributions were left-shifted following Bac_2d_. (*E*) Changes in MFR per neuron. Line indicates no change. (*F*) MFRs were reduced following Bac_2d_. (0.6 ± 0.053 Hz for IGF1R-KO, 0.35 ± 0.053 for KO+Bac). (*G-I)* IGF1R-KO does not affect basal firing properties (same data as in *A-F*). No difference was found in MFR distributions between Ctrl and IGF1R-KO neurons (*G, P* = 0.1), in fraction of spikes participating in bursts (*H, P* = 0.31, 0.23 ± 0.01 for Ctrl, 0.24 ± 0.01 for IGF1R-KO) and in the intra-burst spike frequency (*I, P* = 0.32, 185.1 ± 4.1 Hz for Ctrl, 188.4 ± 4.7 Hz, for IGF1R-KO). Wilcoxon test (*C,F*), Kolmogorov-Smirnov test (*A,D,G*), Mann-Whitney test (*H,I*). Error bars indicate SEM. ns, not significant, \*\**P* < 0.01, \*\*\*\**P* < 0.0001.

We next asked whether cytoCa^2+^ is also homeostatically regulated, and whether IGF1R is necessary for this process. Continuous imaging of cytoCa^2+^ during spontaneous spiking activity in excitatory hippocampal neurons was conducted at baseline, and following 2 days of baclofen perturbation (Fig. S3). The amplitude, frequency and total Ca^2+^ levels (amplitude x frequency) of cytosolic events were quantified for Ctrl and IGF1R-KO neurons. In Ctrl, 2 days of baclofen did not affect cytoCa^2+^ event amplitudes, rate, and subsequently total cytoCa^2+^ (Fig. S3A,C–E). In IGF1R-KO neurons, the amplitude of cytoCa^2+^ events remained unaltered following the perturbation, but their frequency was diminished, thus resulting in lower total cytoCa^2+^ levels (Fig. S3B,F–H) following 2 days of the perturbation. These results further support cytoCa^2+^ as a regulated variable maintained by a homeostatic system and demonstrate that IGF1R deletion impairs this regulation.

Next, we tested if IGF1R is necessary for MFR renormalization to a set-point value for bi-directional changes in activity. We induced chronic hyperactivity in IGF1R-KO neurons by enhancing glutamate spillover by inhibiting glutamate transporters (39). Application of 10 µM TBOA, a competitive glutamate transporter antagonist (40), induced a transient increase in MFR that was gradually renormalized during the following 2 days to the set-point value (Fig. S4). Taken together, these results indicate that IGF1R is essential for upward but not for downward homeostatic restoration of MFRs.

### IGF1R deletion blocks postsynaptic and intrinsic homeostatic plasticity

MFR homeostasis is achieved by intrinsic and synaptic adaptations that act in a negative-feedback manner to counteract disturbances to ongoing activity (12). Is IGF1R necessary for a single but crucial compensatory mechanism, or rather is it an up-stream regulator of distinct compensatory mechanisms? To address this question, we measured several parameters of intrinsic excitability and synaptic strength in excitatory neurons using whole-cell patch clamp. We first tested whether intrinsic excitability is changed after 2 days of baclofen. We elicited action potentials by injecting somatic currents ranging from zero to 600 pA (F-I curves) in the presence of postsynaptic receptor blockers in Ctrl (Fig. 3A,B) and IGF1R-KO (Fig. 3C,D) excitatory neurons. In Ctrl, 2 days of baclofen elicited an increase in firing rate in response to current injections (Fig. 3A,B) and augmented the maximal firing frequency (Fig. S5A). However, this plasticity of intrinsic excitability was lost in IGF1R-KO neurons, as reflected by the lack of change in the F-I curve (Fig. 3C,D) and the maximal firing frequency (Fig. S5A) in response to 2 days of baclofen. Input resistance, spike threshold voltage, spike amplitude and half-width were unaltered by 2 days of baclofen in either Ctrl or IGF1R-KO groups (Fig. S5B–E). It is worth mentioning that IGF1R deletion caused an increase in the input resistance (Fig. S5B) and a decrease in the action potential amplitude (Fig. S5D).

**Figure 3.**
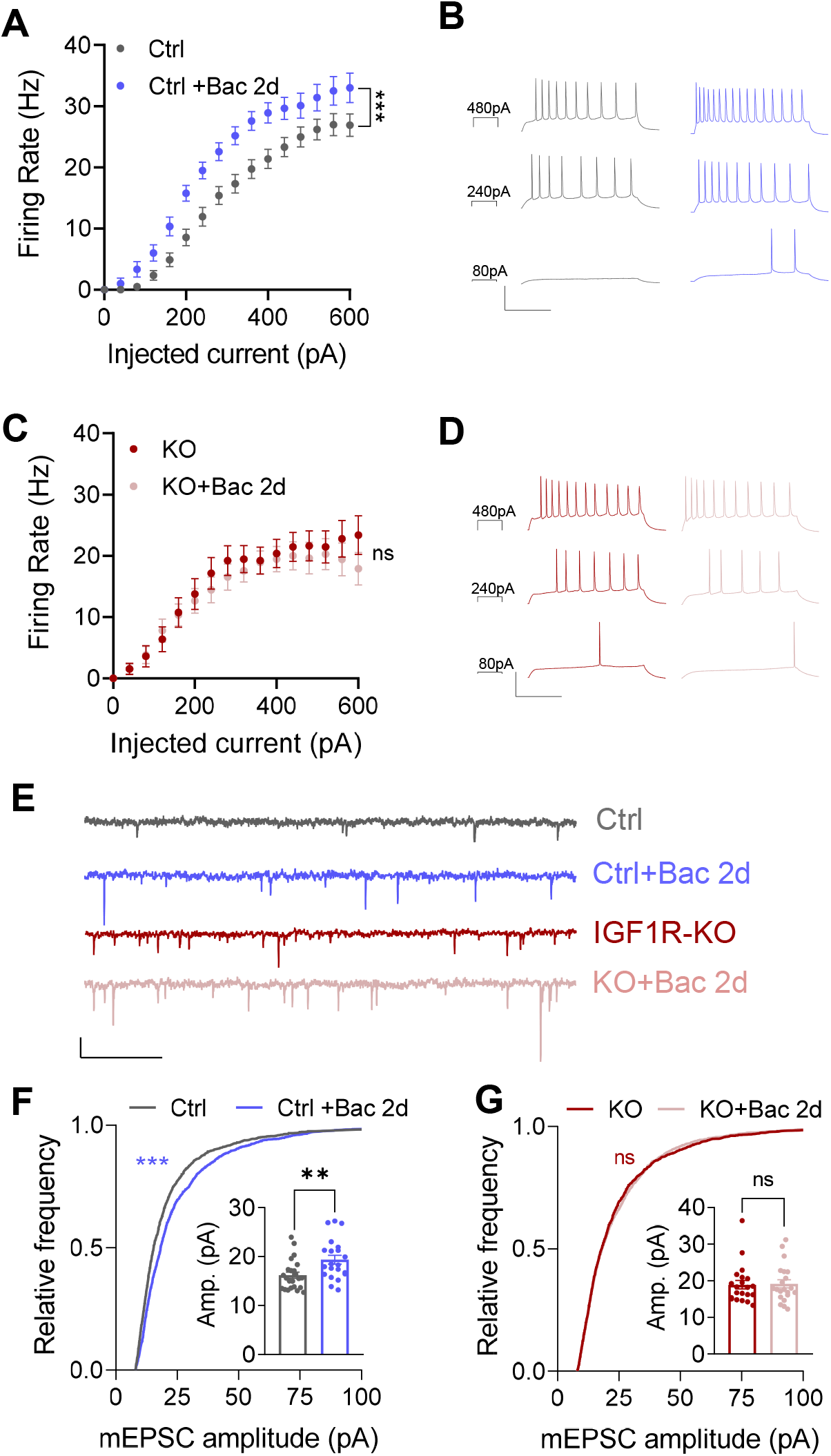
Lack of intrinsic excitability and postsynaptic homeostatic adaptations in IGF1R-KO neurons. (*A*) Increased firing of Ctrl neurons in response to depolarizing currents following Bac_2d_ (n = 29 for Ctrl, n = 24 for Ctrl+Bac). (*B*) Representative traces of Ctrl neurons before (*Black*) and after (Blue) Bac_2d_. (*C*) No change (*P* = 0.56) in firing of IGF1R-KO neurons in response to depolarizing currents following Bac_2d_ (n = 26 for IGF1R-KO, n = 26 for KO+Bac). (*D*) Representative traces of IGF1R-KO neurons before (*Dark red*) and after (*Pink*) Bac_2d_. (*E*) Representative mEPSC recordings of neurons from each group (*scale bars*: 1 sec, 20 pA). (*F*) Cumulative distribution of mEPSC amplitudes in Ctrl was skewed towards larger amplitudes following Bac_2d_ (n = 2061 events from 23 Ctrl neurons and n = 1887 events from 21 Ctrl+Bac neurons). *Inset*: mean of amplitudes is increased following Bac_2d_ (16.2 ± 3.0 pA, n = 23 for Ctrl; 19.37 ± 4.11 pA, n = 21 for KO+Bac). (*G*) Cumulative distribution of mEPSC amplitudes in IGF1R-KO did not change following Bac_2d_ (*P* = 0.87, n = 1860 events from 21 IGF1R-KO neurons, 1890 evets from 21 KO+Bac neurons). *Inset*: mean of mEPSC amplitudes did not change (*P* = 0.87) following Bac_2d_ (18.92 ± 1.15 pA, n = 21 for IGF1R-KO; 19.17± 1.12 pA, n = 21 for KO+Bac. Two-way ANOVA mixed-effects analysis (*A,C*), Kolmogorov-Smirnov and Mann-Whitney tests (*F,G*). Error bars indicate SEM. ns, not significant, \*\**P* < 0.01, \*\*\**P* < 0.001.

Next, we tested whether IGF1R deletion alters presynaptic and postsynaptic adaptations to inactivity. Indeed, 2 days of baclofen elicited a marked increase in the amplitude of miniature excitatory post-synaptic currents (mEPSCs) in Ctrl neurons (Fig. 3E,F), whereas this homeostatic postsynaptic plasticity was lost in IGF1R-KO, reflected by the lack of baclofen effect on the mEPSC amplitude (Fig. 3E,G). On the other hand, the mEPSC frequency was increased in both groups (Fig. S6), indicating that presynaptic homeostatic mechanisms in excitatory neurons are independent of IGF1R. Taken together, our results demonstrate that deletion of IGF1Rs impaired both postsynaptic and intrinsic homeostatic mechanisms that may contribute to the failure of MFR homeostatic recovery at the network level.

### IGF1R deletion suppresses spike-to-mitoCa^2+^ transfer function

How changes in firing rates and patterns are translated into physiological error signals within neurons remains unknown. Somatic cytoCa^2+^ has long been assumed to be a regulated variable through which neurons sense changes in spiking activity, and by doing so are able to correct deviations from MFR set points (18, 19). Mitochondria are known to buffer Ca^2+^ upon neuronal activation (41), and thus may serve as a sensor of activity-induced changes in cytoCa^2+^ (17). Given that a fraction of IGF1Rs is localized in mitochondria, we asked how IGF1Rs regulate activity-dependent cytoCa^2+^ and mitoCa^2+^ in soma of excitatory neurons. To address this question, we measured spike-to-cytoCa^2+^ and cyto-to-mitoCa^2+^ transfer functions for different patterns of spikes in Ctrl versus IGF1R-KO neurons. We conducted simultaneous dual-color imaging of cytosolic and mitochondrial Ca^2+^ dynamics by using jRGECO1a and 4mt-GCaMP8m, respectively (Fig. 4A). To identify excitatory neurons, we used 4mt-GCaMP8m under the CaMKIIα promoter. Neurons were stimulated by a single spike and spike bursts comprising of 3, 5 and 10 spikes at 50 Hz, while spontaneous firing was blocked. Under these conditions, spike-to-cytoCa^2+^ transfer functions did not differ between Ctrl and IGF1R-KO neurons (Fig. 4B,D). Conversely, IGF1R deletion caused a significant reduction in spike-to-mitoCa^2+^ transfer function (Fig. 4C,E). Importantly, mitoCa^2+^ transients were activated only by spike bursts, but not by single spikes. Hence, IGF1R deletion diminished Ca^2+^ uptake by mitochondria evoked by spike bursts only. These changes resulted in a right shift of the cytoCa^2+^-to-mitoCa^2+^ transfer function (Fig. 4F).

**Figure 4.**
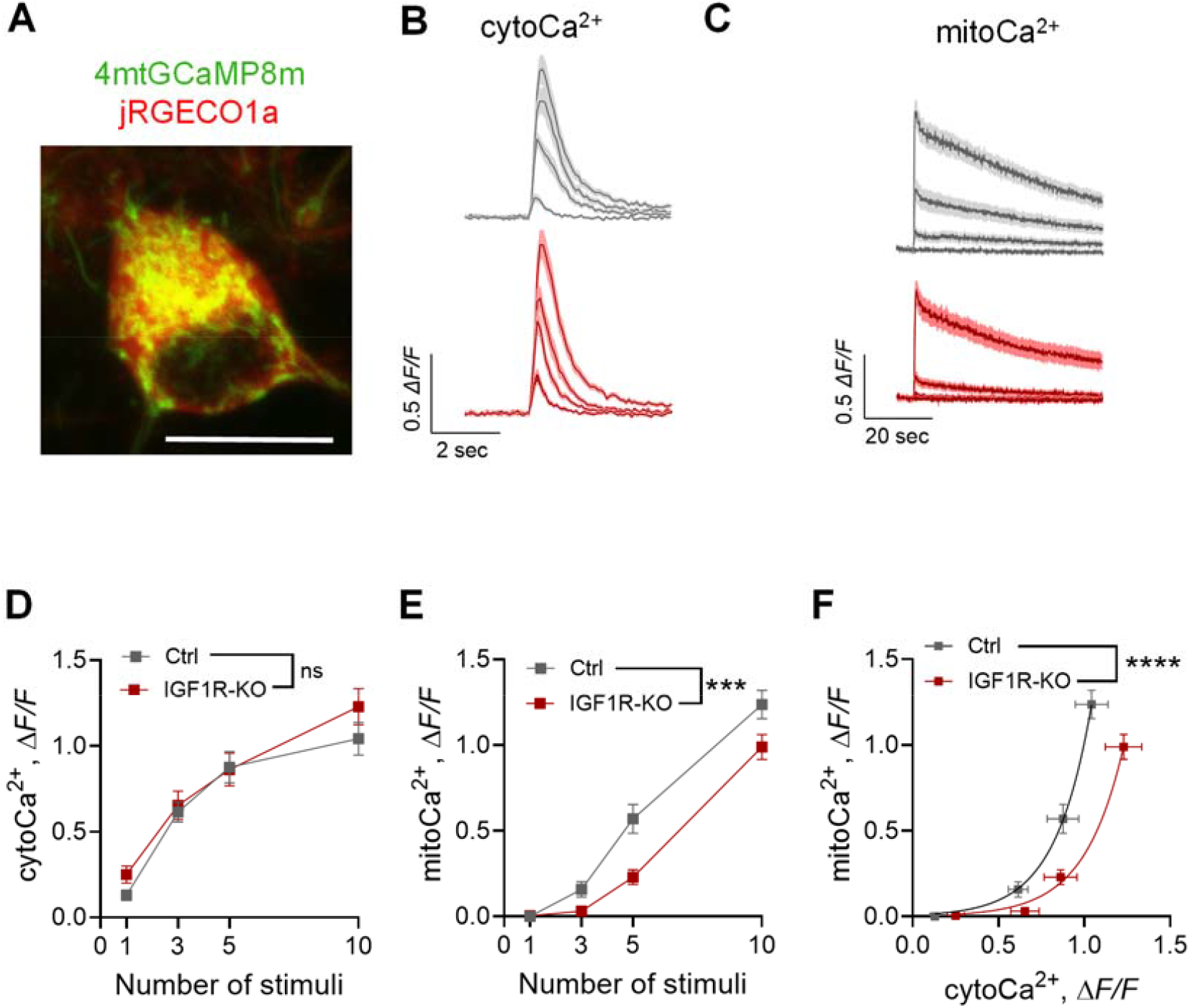
IGF1R deletion decreases somatic mitoCa^2+^ and cytoCa^2+^-to-mitoCa^2+^ coupling evoked by spike bursts. (*A*) An excitatory neuron expressing cytosolic jRGCECO1a and mitochondrial GCaMP8m. Scale bar: 10 µm. (*B*) Mean (*dark line*) and SEM of cytoCa^2+^ traces for a single spike and spike bursts composed of 3,5,10 stimuli at 50 Hz. Scale bars: 0.5 ΔF/F, 2 sec. (*C*) Mean (*dark line*) and SEM of mitoCa^2+^ traces for a single spike and spike bursts composed of 3,5,and 10 stimuli at 50 Hz. *Scale bars*: 0.5 ΔF/F, 20 sec. (*D*) CytoCa^2+^ responses were not changed (*P* = 0.42) by deletion of IGF1R (n = 40 - 42 for Ctrl, n = 32 - 37 for IGF1R-KO). (*E*) MitoCa^2+^ transients evoked by spike bursts were reduced in IGF1R-KO neurons (n = 40 - 42 for Ctrl, n = 32 - 37 for IGF1R-KO). (*F*) CytoCa^2+^-to-mitoCa^2+^ coupling was reduced in IGF1R-KO (n = 165 events for Ctrl, n = 142 events for IGF1R-KO). Squares are mean of cytoCa^2+^ events from (*D*) and mean of mitoCa^2+^ events from (*E*). Two-way ANOVA mixed effects analysis (*D,E*), Least square exponential regression. Extra sum-of-squares F test (*F*). Error bars indicate SEM. ns, not significant, \*\*\**P* < 0.001, \*\*\*\**P* < 0.0001.

Given the central role of mitochondria in ATP production, we tested whether IGF1R deletion suppressed ATP homeostasis. We used the cytosolic FRET sensor ATeam (42) to estimate somatic ATP levels in Ctrl and IGF1R-KO hippocampal neurons. Our results show no difference in FRET efficiency by IGF1R deletion during spontaneous activity (Fig. S7). Taken together, these data suggest that IGF1R-KO specifically impairs somatic Ca^2+^ uptake by mitochondria, without affecting cytoCa^2+^ and ATP levels. The decreased mitoCa^2+^ transients elicited by burst-induced cytoCa^2+^ events, ultimately lead to a weakening of cytoCa^2+^-to-mitoCa^2+^ coupling.

### Rescue of mitoCa^2+^ and upward MFR homeostatic response by MCUc

Given that MCUc is the primary Ca^2+^ source into mitochondria (37), reduction in mitoCa^2+^ by IGF1R deletion may result from downregulation of the MCUc subunits. To test this hypothesis, we measured the mRNA expression levels of the pore-forming subunit MCU, as well as the Ca^2+^-sensing subunits MICU1, MICU2 and MICU3. We found that IGF1R deletion caused a downregulation in the expression level of MCU and the brain-specific (43) MICU3 subunits, while it did not affect the expression levels of MICU1 and MICU2 subunits (Fig. 5A). Thus, IGF1R affects MCU and MICU3 expression at the transcriptional level.

**Figure 5.**
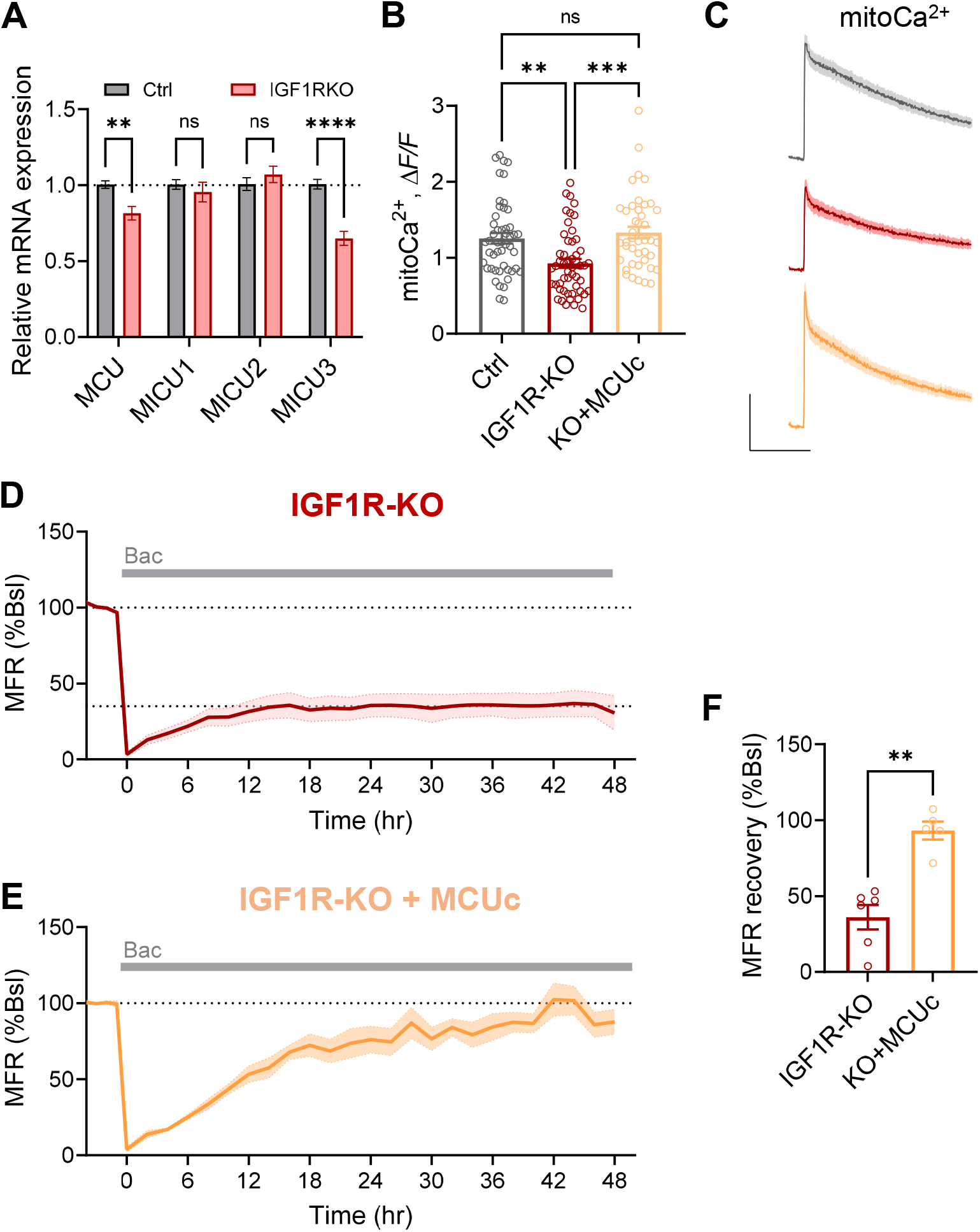
Over-expression of MCUc rescues mitoCa^2+^ and MFR homeostasis in the absence of IGF1R. (*A*) mRNA expression of MCUc proteins in IGF1R-KO cultures relative to Ctrl: The mRNA of MCU and MICU3 was reduced in IGF1R-KO culture (0.815 ± 0.045 for MCU; 0.649 ± 0.046 for MICU3), and did not change in MICU1 and MICU2 (0.954 ± 0.064, *P* = 0.925 for MICU1; 1.070 ± 0.054, *P* = 0.832 for MICU2, n = 15 for IGF1R-KO; n = 13 for Ctrl. Samples from 3 experiments for MCU and MICU3, samples from 2 experiments (n = 9) for MICU1 and MICU2). (B) MitoCa^2+^ in response to 10 stimuli at 50 Hz is reduced by IGF1R deletion, and is rescued by over-expression of MCUc in IGF1R-KO neurons (*P* > 0.99, 1.25 ± 0.07, n = 49 for Ctrl; 0.93 ± 0.06, n = 52 for IGF1R-KO, 1.33 ± 0.08, n = 42 for KO+MCUc). (*C*) Mean (*dark line*) and SEM of mitoCa^2+^ event traces in (*B*). (*D*) Impaired MFR response to chronic inactivity induced by 10 µM baclofen in IG1R-KO cultures expressing mCherry, normalized to the baseline MFR (6 experiments, 291 channels). (*E*) Restoration of a normal MFR homeostatic response in IGF1R-KO+MCUc, normalized to the baseline MFR (5 experiments, 252 channels). (*F*) Summary (same data as *D-E*) shows restoration of MFR recovery following Bac_2d_ by MCUc in IGF1R-KO (93.1 ± 5.9% of baseline for IGF1R-KO+MCUc compared to 36.1 ± 8.0 % for IGF1R-KO+mCherry). Each circle is the mean of the last four time-points of each experiment in (*D*) and (*E*). Two-way ANOVA with Sidak’s multiple comparison test (*A*), Kruskal-Wallis test with Dunn’s correction for multiple comparisons (*B*), Mann-Whitney test (*F*). Error bars indicate SEM. ns, not significant, \*\**P* < 0.01, \*\*\**P* < 0.001.

Next, we tested whether IGF1R is colocalized in mitochondria with MCUc, using 2-color stimulated emission depletion (STED) microscopy (Fig. S8). This revealed a partial colocalization of the four MCUc components and IGF1R (Fig. S8C,C’). Quantitative analysis indicated that the colocalization is statistically significant, with the IGF1R signal in the MCUc spots being significantly higher than elsewhere (Fig. S8D). On average, ~30% of IGF1Rs in soma are localized with the *bona fide* mitochondrial marker TOM20 (Fig. S8E). Taken together, these results suggest a partial, but significant colocalization of IGF1R with MCUc complexes.

To directly test whether reduction in mitoCa^2+^ is the primary cause of impaired MFR homeostasis in IGF1R-KOs, we tested whether ectopic overexpression of MCUc may rescue mitoCa^2+^ and MFR homeostasis. Our results show that overexpression of MCU, together with its regulatory subunits (MICU1 and MICU3), led to an increase in mitoCa^2+^ transients evoked by spike bursts in IGF1R-KOs, rescuing it to the control level (Fig. 5B,C). Next, we tested if the rescue of mitoCa^2+^ in IGF1R-KO neurons is sufficient to restore MFR homeostasis in response to inactivity. While MFR recovered only to 36% of the baseline in IGF1R-KO networks over-expressing mCherry (Fig. 5E,F), over-expression of MCUc markedly increased the compensatory response of IGF1R-KO networks to 93% of the baseline (Fig. 5D,F). As MCUc rescues both mitoCa^2+^ and MFR homeostasis in response to inactivity, these results strongly suggest that IGF1R deletion impairs MFR homeostatic recovery from inactivity by reducing mitoCa^2+^ uptake during spike bursts.

## Discussion

Long-term stability of ongoing spiking dynamics is crucial for neural circuits’ functions (44–48). Although a large number of fine-tuned parameters regulating synaptic and intrinsic membrane properties can generate similar firing properties (49, 50), the cellular and molecular design underlying MFR homeostasis in central neural networks remains largely unknown. Here, we identify a novel role of IGF1Rs, known regulators of brain development (21), proteostasis (51, 52) and lifespan (23, 24, 26), in homeostasis of neural network activity. Our results provide converging evidence on the necessity of evolutionary-conserved IGF1R signaling in the stabilization of firing rate distributions at the population level in hippocampal networks. These results are important for several reasons. First, they demonstrate that IGF1Rs are dispensable in regulating MFR set points during spontaneous neuronal activity, but are critical for the homeostatic compensation of MFR to inactivity. Second, they reveal a role of IGF1R in regulating temporal Ca^2+^ filtering via MCUc of neuronal mitochondria during periods of spike bursts. Third, they show that MCUc can restore upward MFR homeostasis in IGF1R-deficient networks. Finally, they point to a critical role of mitochondria in regulating the integrated homeostatic response at the network level.

### Mitochondria as high pass filters in central neurons

Our results indicate that neurons are extremely unreliable at transferring information encoded by single spikes to somatic mitochondria. The coupling of mitochondria-to-cytosolic Ca^2+^ is non-linear, showing almost complete uncoupling during periods of low-frequency, single spikes. However, spike bursts, known to play an important role in synaptic plasticity and information processing (53), are reliably signaled to mitochondria by activating mitoCa^2+^ influx via MCUc. Thus, neuronal mitochondria can be viewed as filters that transmit bursts, but filter out single spikes. Our results demonstrate that these filter properties are regulated by IGF1Rs. Whether MCUc is regulated via an intracellular messenger activated by plasma membrane IGF1Rs or via the mitochondria-residing IGF1R fraction remains to be uncovered by future studies. Irrespective of the precise mechanism, our results point to the IGF1R / MCUc signaling pathway as a key filter of intracellular Ca^2+^ dynamics during periods of correlated activity that may be important for the induction of homeostatic plasticity mechanisms. While our conclusions are based on cultured neurons, a recent *in vivo* study in the cortex of awake mice showed a positive relationship between mitoCa^2+^ and cytoCa^2+^ peak amplitudes, wherein large cytoCa^2+^ events are likely to be evoked by spike bursts (54). Therefore, unreliable spike-to-mitoCa^2+^ coupling during single spikes appears to be a universal property of somatic mitochondria in central neurons. Given that duration of mitoCa^2+^ events is ~8 times longer than cytoCa^2+^, mitochondria perform amplification and high-pass filtering of burst-evoked cytoCa^2+^ signals. Thus, MCUc does not only sense cytoCa^2+^ to control the threshold and gain as has been previously proposed (55), but also controls information content transferred from cytoplasmic membrane potential to mitochondria.

Previous findings suggest that L-type voltage-gated calcium channels are necessary for homeostatic plasticity of intrinsic excitability (56, 57), while an endoplasmic reticulum (ER) Ca^2+^ sensor (58) and T-type calcium channels (59) are required for synaptic homeostatic plasticity. Interestingly, L-type calcium channels have been demonstrated to play a critical role in transcription, while mitochondria and ER buffer Ca^2+^ entering via N-type calcium channels, limiting their involvement in transcriptional regulation (60). Thus, a decrease of Ca^2+^ entry to the mitochondria, induced by loss of IGF1Rs, may enhance excitation-transcription coupling. How IGF1R / MCUc signaling regulates mitochondria-to-nucleus Ca^2+^ coupling and transcriptional programs related to induction of upward MFR homeostatic response remains a challenge for future research.

### IGF1R and MFR homeostasis

The network’s ability to yield the same output despite different molecular compositions, called degeneracy, is proposed to be a ubiquitous biological property at all levels of organization (61). However, homeostatic regulation may fail when one of the core homeostatic machinery components becomes dysfunctional (16). Here, we show that IGF1R deficiency limits upward firing rate homeostasis by suppressing mitoCa^2+^-to-cytoCa^2+^ coupling via MCUc. Thus, in addition to cytoCa^2+^, mitoCa^2+^-to-cytoCa^2+^ coupling may play an important role in MFR homeostasis.

It has been recently demonstrated that the loss of Shank3, implicated in autism spectrum disorders, impairs upward MFR homeostasis and ocular dominance plasticity in the V1m cortex (62). Interestingly, injection of IGF-1 prevents effects of monocular deprivation on ocular dominance plasticity in the V1 cortex (28). Moreover, IGF1 restored deficits in excitatory synaptic transmission in neurons with reduced Shank3 expression from 22q13 deletion syndrome patients (63). However, whether deficiency in mitoCa^2+^ evoked by spike bursts also limits upward MFR homeostasis in visual cortex and hippocampus *in vivo* is unknown. Whether common homeostatic mechanisms maintain stability of distinct neural circuits serving different functions remains an attractive question for future studies.

We hypothesize that maintaining a delicate balance of IGF1R signaling is critical for normal brain functioning. On the one hand, some brain disorders are associated with down-regulation of IGF1. For example, *Mecp2* mutant mice, a model of Rett syndrome, exhibit decreased levels of serum IGF1 (64). A treatment of *Mecp2* mutant mice with systemic IGF1 restored Rett syndrome-like symptoms, including synaptic and cognitive deficits (64, 65). As *Mecp2* deletion impairs homeostatic synaptic scaling (66, 67) and excitation-to-inhibition balance (68), future studies are needed to test if IGF1 can restore the symptoms by rescuing homeostatic failures. On the other hand, a decrease in IGF1R signaling protects from amyloid-β-mediated pathology as well as from synaptic, neuronal and cognitive deficits in Alzheimer’s disease mouse models (34, 51, 69, 70). Moreover, inhibition of MCU decreases mitoCa^2+^ overload in cortical neurons of Alzheimer’s disease model mice (71). Reduced IGF1R / MCUc signaling may be neuroprotective by suppressing disease-associated hyperactivity of hippocampal synapses (34) and by restricting upward MFR homeostasis. It is noteworthy that the transcription factor REST regulates downward MFR homeostasis (72) and extends lifespan (73). It remains to be seen whether a critical role of IGF1R in development and ageing is rooted in its capacity to regulate firing rate homeostasis of central neural circuits.

## Methods

### Primary hippocampal culture preparation

Hippocampi were dissected from IGF1R^fl/fl^ (74) pups (both sexes) at P0-1in ice cold Leibovitz L-15 medium. Tissue was washed 3 times with Hank’s balance salt solution (HBSS). Chemical dissociation of cells was done using digestion solution (137 mM NaCl, 5 mM KCl, 7 mM Na_2_HPO_4_, 25 mM HEPES, 2 mg/mL trypsin, 0.5 mg/mL DNase) for 10 min in an incubator. Solution was replaced by HBSS supplemented with 20% FBS to inactivate trypsin and once again with only HBSS. Cells were then mechanically dissociated in HBSS supplemented with 13 mM MgSO_4_ and 0.5 mg/mL DNase by titration with fire-polished pipettes of decreasing diameter. Sedimentation of cells was accomplished by centrifugation at 1000 r*cf* for 10 min at 4°C. After the removal of the supernatant, cells were re-suspended with plating medium (MEM supplemented with 10% FBS, 32.7 mM glucose, 25 mg/mL insulin, 2 mM Glutamax, 0.1 mg/mL transferrin, 0.1% SM1) and then plated on matrigel-coated glass coverslips, glass-bottom 24-wells or MEA plates. Half of the serum medium was replaced with feeding medium the day after (MEM supplemented with 32.7 mM glucose, 2 mM Glutamax, 3 μM ARA-C, 0.1 mg/mL transferrin, 2% SM1). Afterwards, half of the medium was replaced with fresh feeding medium every 3-4 days for three additional times. For all electrophysiology and live cell imaging, cultures were infected after 5-6 days *in-vitro* (DIV) with Cre+ or Cre-AAVs (see next section). The experiments were performed in cultures after 14 – 21 DIV.

### Plasmids

For CBAP-Cre-P2a-Cerulean and CBAP-P2a-Cerulean, cDNA encoding for Cre was obtained from K.Villa. AAV2-CBAP plasmid was obtained from Daniel Gitler (BGU, Israel). Cre-P2a-Cerulean or P2a-Cerulean was inserted into AAV2-CBAP by Gibson assembly between NotI and HindIII or EcoRI and HindIII respectively. AAV-hSynI-jRGECO1a was prepared by cloning jRGECO1a (pGP-CMV-NES-jRGECO1a, Addgene plasmid # 61563) between BglII and NotI sites of AAV2-hSynI. AAV2-hSyn-AT1.03 NL was a gift from Daniel Gitler. AAV2-CaMII2a-4mtsGCaMP8m was based on the plasmid on pAAV-CaMKIIa-GCaMP6s-p2A-nls_dTomato (Addgene #51086). Kozak-4MTS fragment was synthesized by GenScript and cloned into the intermediate plasmid-AAV2-hSyn-linker-jGCaMP8m. The latter was constructed by cloning linker-jGCaMP8m (Addgene #162372) into AAV2-hSyn1.The entire cassette Kozak-4MTS-jGCaMP8m was cloned into pAAV-CaMKIIa between SpeI and EcoRI sites. For the production of CBAP-mMCU-mCherry, CBAP-mMICU1-mCherry and CBAP-MICU3-mCherry, mMCU (NM_001033259.4) and mMICU1 (NM_144822.3) were synthesized by GenScript. mMICU3 (NM_030110.2) was synthesized by Genwiz. Initially, P2a-mCherry was cloned into AAV2-CBAP, then mMcu, MICU1 or MICU3 was cloned upstream of P2a.

### Electrophysiology

#### MEA data acquisition and analysis

Cells were grown on MEA plates [Multi Channel Systems (MCS), 120MEA200/30iR-Ti] containing 120 titanium nitride (TiN) electrodes with 4 internal reference electrodes. Each electrode’s diameter is 30 μm and they are spaced on a 12×12 grid (24 spaces in the 4 corners did not contain electrodes), spaced 200 μm apart. Data was recorded by either a MEA2100-System (MCS) with a chamber that maintained 37°C and 5% CO2, or a MEA1200-mini-system (MCS) that was constantly placed inside an incubator. Raw data was collected at 10 kHz, with a hardware high-pass filter of 1 Hz and an upper cut off of 3.3 kHz for the MEA2100-system and 3.5 kHz for the MEA2100-mini-system.

Using MCS data analyzer software offline, raw data were filtered by a Butterworth 2^nd^ order high-pass filter at 200 Hz. Spikes were then detected by a fixed threshold of 6 SD. To reduce processing and analysis time, each hour was represented by 20 min of recording, which was previously shown to reliably represent the MFR of the full hour (4). For spike sorting, Plexon offline-sorter V3 (Plexon inc. USA) was used. Principle component analysis was carried out on a 2-D or 3-D space.

Distinct clusters were manually selected, and an automatic template sorting was done on the rest. Clusters’ stability throughout the recording was manually inspected. Only clusters that fulfilled the following requirements were considered units and used for the analysis: (1) No spikes in refractory periods. (2) The clusters were well defined during the entire experiment. (3) There were no sudden jumps in cluster location on PC axes. The rest of the analysis was carried out using a custom-written MATLAB (Mathworks) as described in (4). For detection of bursts at the single-unit level, bursts were defined as 2 or more spikes at a minimum of 20 Hz based on the code we previously published (4). Our previous analysis shows that the results are robust over a wide range of parameters (4).

#### Patch clamp whole-cell recordings and analysis

Experiments were performed at room temperature in a RC-26G recording chamber (Warner instrument LCC, USA) on the stage of FV300 inverted confocal microscope (Olympus, Japan) using a Multiclamp 700B amplifier and a Digidata 1440A digitizer (Molecular devices, LLC, USA). All experiments were carried out using extracellular Tyrode solution containing (in mM): NaCl, 145; KCl, 3; glucose, 15; HEPES, 10; MgCl_2_, 1.2; CaCl_2_, 1.2; pH adjusted to 7.4 with NaOH, and intracellular solution containing (in mM): K-gluconate 120; KCl 10; HEPESs 10; Na-phosphocreatine 10; ATP-Na_2_ 4; GTP-Na 0.3; MgCl_2_ 0.5. Intracellular solution was supplemented with 20 μM Alexa fluor 488 for dendritic spines imaging. For intrinsic excitability, synaptic blockers were used (in μM): 10 CNQX, 50 AP-5, and 10 gabazine. For mEPSCs recordings, tetrodotoxin (1 μM), AP-5 (50 μM), and gabazine (10 μM) were added to Tyrode’s solution. Intrinsic excitability protocols: small DC current was injected in current-clamp mode to maintain membrane potential at −65 mV. For I-F curve: positive currents from 40 to 600 pA were injected in 40 pA increments for 500 ms. Input resistance (R_in_) was measured by calculating the slope of the voltage change in response to increasing current injections from −80 to +20 mV in 20 mV increments. For single AP measurements, 2 ms currents were injected at 40 pA increments. For mEPSCs recordings, neurons were voltage-clamped at −65 mV. Neurons were excluded from the analysis if no dendritic spines were observed, serial resistance was > 15 MΩ, serial resistance changed by >20% during recording, or if R_in_ was < 75 MΩ. Signals were recorded <u>at 10</u> kHz, and low-pass filtered with Bessel filter 2 kHz. Electrophysiological data were analyzed using pClamp (Molecular Devices LLC, USA) and MiniAnalysis (Synaptosoft, Decatur, Georgia, USA) for mEPSC. For distributions of mEPSCs, 90 first events were taken from each neuron to prevent over-representation of high-frequency neurons.

### Confocal live cell imaging

#### Evoked cytosolic and mitochondrial calcium imaging in neuronal soma

Experiments were performed on a FV-1000 (Olympus, Japan) system using a QR/RC-47FSLP chamber with a TC-324C temperature controller (Warner Instruments LLC, USA) at 33-34°C. In-chamber temperature stability was verified between coverslips with a Newtron TM-5005 thermometer. Coverslips were placed in Tyrode’s solution that contained (in mM): NaCl, 145; KCl, 3; glucose, 15; HEPES, 10; MgCl_2_, 1.2; CaCl_2_, 1.2; pH adjusted to 7.4 with NaOH at 34°C. Synaptic blockers were used to prevent recurrent activity from stimulation (in uM): 10 CNQX, 50 AP-5. Cultures were infected with AAV1/2-hSyn-jRGECO1a and AAV1/2-CaMKIIα-4mt-GCaMP8m, so that only excitatory neurons expressed both Ca^+2^ sensors. Only cells with mitochondrial response to a 10-stimuli burst at 50 Hz were considered. Infection with AAV1/2-CBAP-Cre-Cerulean or AAV1/2-CBAP-Cerulean was verified with a 440 nm laser. Imaging was done with a 60X lens and X1.5 digital magnification at ~12.5 frames per second with 488 nm and 561 nm lasers, and emission spectra of 505 – 540 nm and 575 – 675 nm for GCaMP8m and jRGECO1a, respectively. Field stimulation was given using a SIU-102 stimulation unit (Warner Instruments LLC, USA) connected to an Axon Digidata 1440A digitizer (Molecular devices, LLC, USA). Each imaged neuron was stimulated by a single stimulus and bursts of 3, 5 and 10 stimuli at 50 Hz. Some neurons had more than one clear response to 3, 5 or 10 stimuli and thus excluded from analysis. Analysis were done using ImageJ.

#### Spontaneous cytosolic calcium in neuronal soma

Experiments were performed on a FV-1000 (Olympus, Japan) system. Cultures were plated on #0 glass bottom 24 well (Cellvis, USA) and imaged in a heated (34-35°C) Stage-Top Incubator System TC, connected to a CU-501 temperature controller and a humidifier delivering 5% CO2 air (Live Cell Instruments, Republic of Korea). Imaging parameters were the same as for the evoked imaging. Each neuron was recorded 3 times for 3 minutes, with a 3-minute interval to prevent bleaching and phototoxicity. Analysis was done using ImageJ (for extraction of florescence intensity over time) and a custom-written MATLAB script (for dF/F calculation and event identification and quantification).

### Real-time PCR

RNA was extracted using the TRI Reagent, according to the manufacturer’s instructions (Sigma-Aldrich). Equal amount of mRNA was reverse-transcribed to cDNA with Superscript III reverse transcriptase (Invitrogen, cat. No: 18080-051). Real-time qPCR was performed with Sybr mix (Applied Biosystems). Reactions were run in triplicate in a StepOnePlus real-time PCR system (Applied Biosystems). mRNA abundance was calculated by means of the comparative cycle threshold (Ct) method following the manufacturer’s guidelines. mRNA expression level was normalized to the average expression of B*2m* and *Polr2a*.

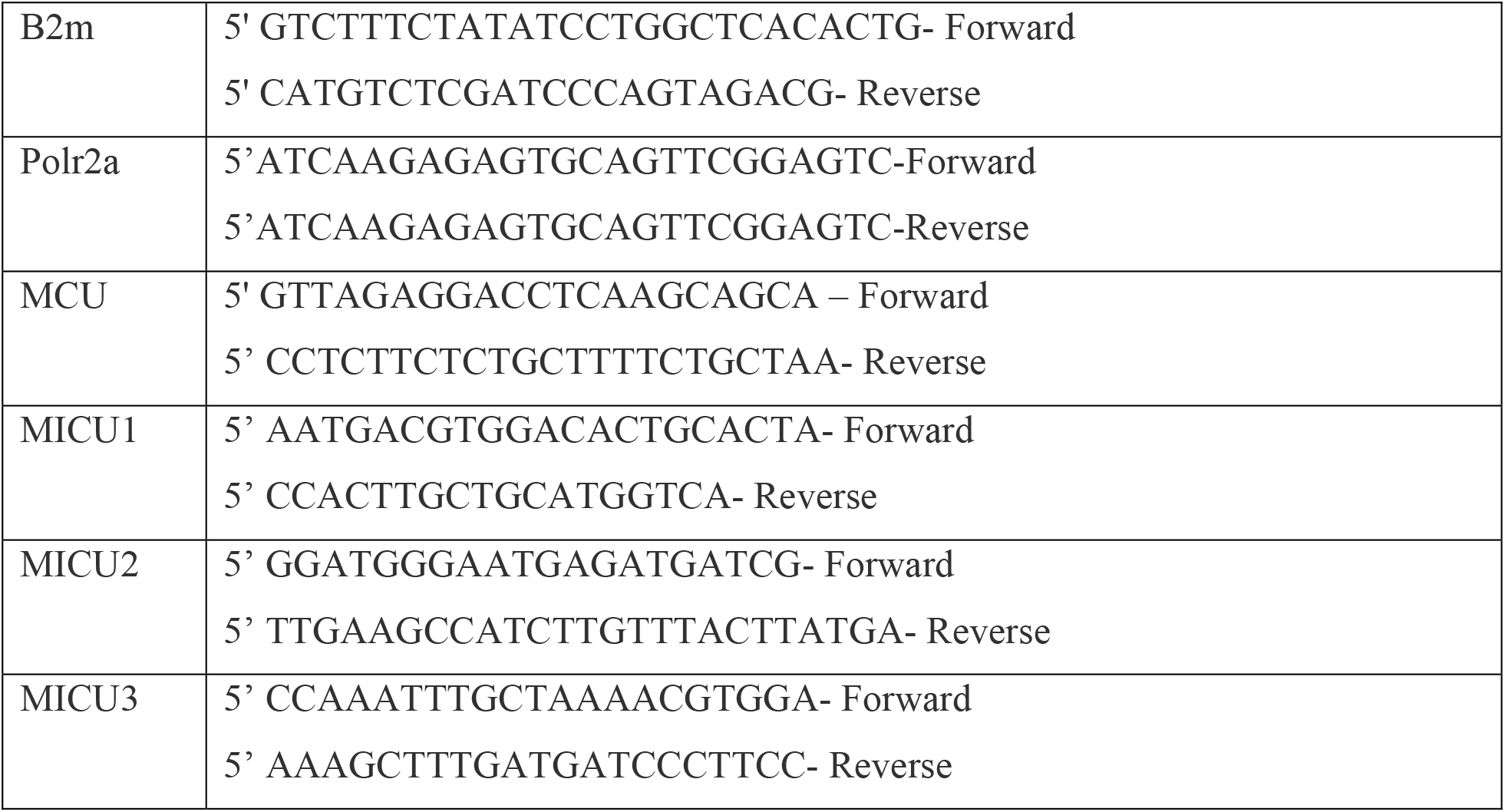

### AAV vector production

AAV vector production was carried out in 293T cells. Cells were transfected 24 h after seeding with helper plasmids encoding AAV rep, cap and plasmid for the rAAV cassette expressing the relevant DNA. Cells were harvested 72 h after transfection, cells pellet was resuspended in lysis solution (150 mM NaCl, 50 mM Tris-HCl, pH 8.5), 1mL of lysis buffer per 150 mm dish. Cells were lysed by a few freeze-thaw cycles. The obtained crude lysate was treated with 50 U benzonase (Sigma, E1014) per 1 mL of lysate at 37°C for 1.5 h to degrade genomic DNA. Cell debris were pelleted by centrifugation at 3000*g* for 15 min 4°C. Supernatant contains crude virus was filtered through a 0.45 um filter and stored at 4°C.

### Statistical analysis

Experiments were repeated at least three times (three independent cultures) for each group. Data are shown as mean ± standard error of the mean (SEM). All statistical analyses were carried out using GraphPad Prism 8.0 (GraphPad software, USA), and the specific tests of each figure, along with their *P*-values and total number of cells/repetitions per experimental group (n/N) are specified in the figure legends.

## Acknowledgments

We thank Tim O’Leary and Lee Susman for comments on the manuscript and all members of the Slutsky Lab for discussions. This work was supported by research grants to I.S. from the European Research Council (724866), the Israel Science Foundation (1663/18), Rosetrees Trust (A2590), the Volkswagen Foundation and the Ministry of Science and Culture of Lower Saxony grant (with Dr. Silvio O. Rizzoli), the Deutsche Forschungsgemeinschaft (440813539 with Dr. Silvio Rizzoli), and BIRAX Regenerative Medicine Initiative (with Dr. Tara Keck). S.A. and M.H. received research funds from INSERM (Institut National de la Santé et de la Recherche Médicale) and Sorbonne University. I.S. is grateful to Sheila and Denis Cohen Charitable Trust and Rosetrees Trust of the UK for their support. M.K. is grateful to Sagol School of Neuroscience for the award of doctoral fellowship. This work was performed in partial fulfillment of the requirements for a Ph.D. degree by Maxim Katsenelson at the Sackler Faculty of Medicine and Sagol School of Neuroscience, Tel Aviv University, Israel.

## Supplementary Figures

**Figure S1.**
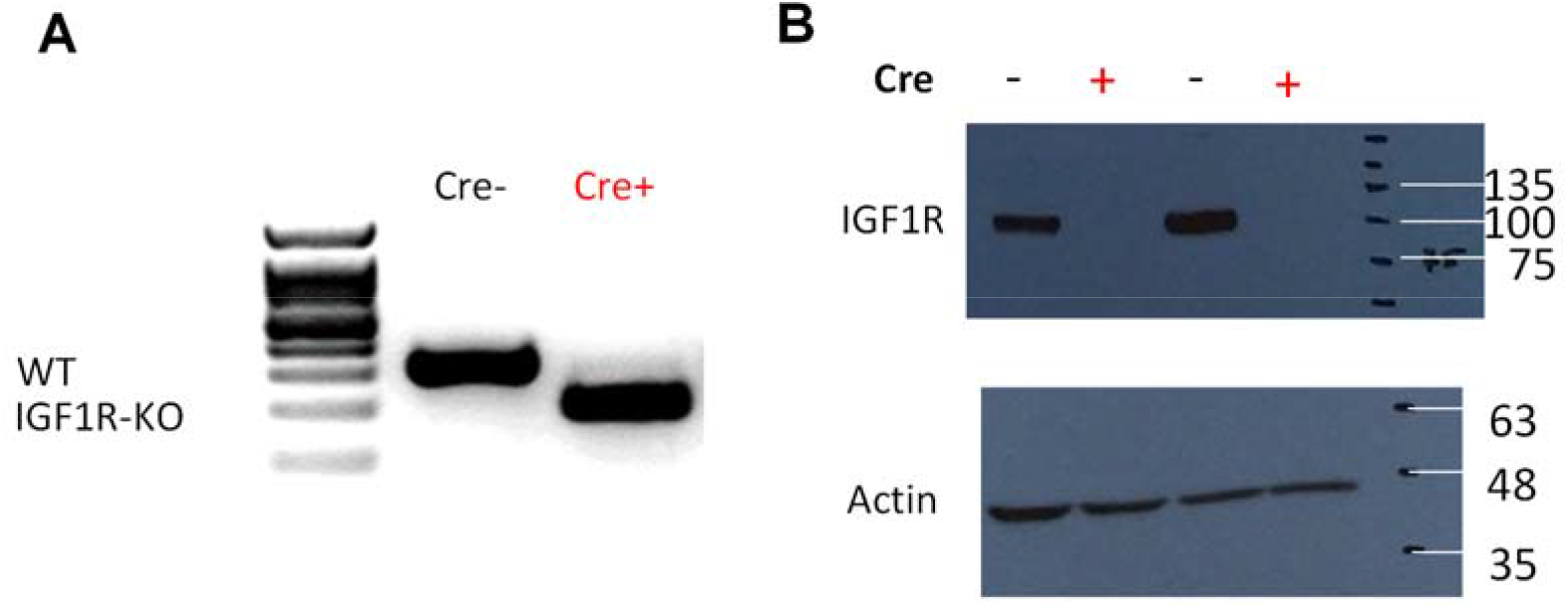
Complete excision of IGF1R gene and elimination of IGF1R protein in Cre-infected IGF1R-fl/fl cultures. Cultures were collected to produce DNA and Protein. (*A*) IGF1R ^flox/flox^ vs. excised IGF1R genes in IGF1R-KO and Ctrl cultures. (*B*) Western-blot of IGF1R in IGF1R-KO and Ctrl cultures.

**Figure S2.**
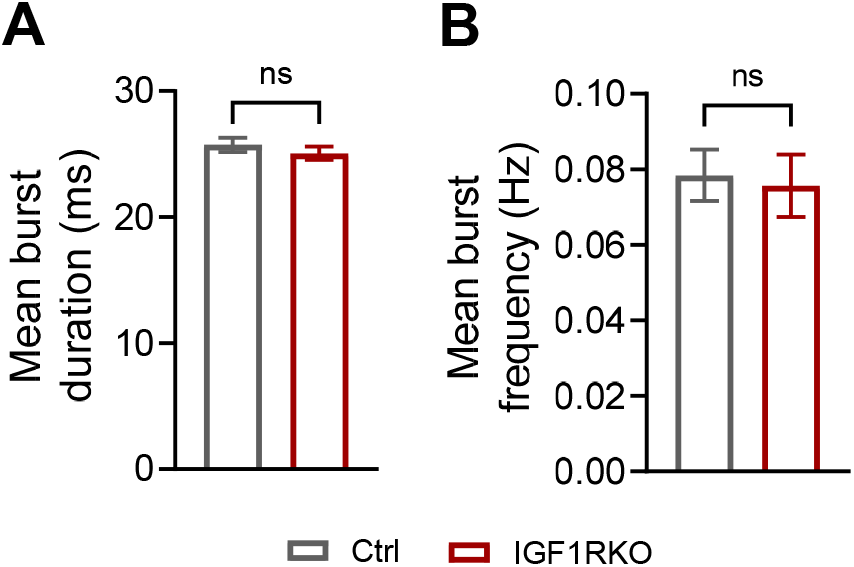
Deleltion of IGF1R does not change the majority of basal spiking-patterns parameters, but prevents their adaptations to inactivity. (*A-B*) No difference between Ctrl and IGF1R-KO in the mean burst duration (*A, P* = 0.98, 25.8 ± 0.6, n = 513 for Ctrl; 25.1 ± 0.5, n = 342 for IGF1R-KO) and in the frequency of bursts (B, *P*=0.56, 0.078 ± 0.007, n=519 for Ctrl; 0.076 ± 0.008, n = 347 for IGF1R-KO). Mann-Whitney test. Error bars indicate SEM. ns, not significant.

**Figure S3.**
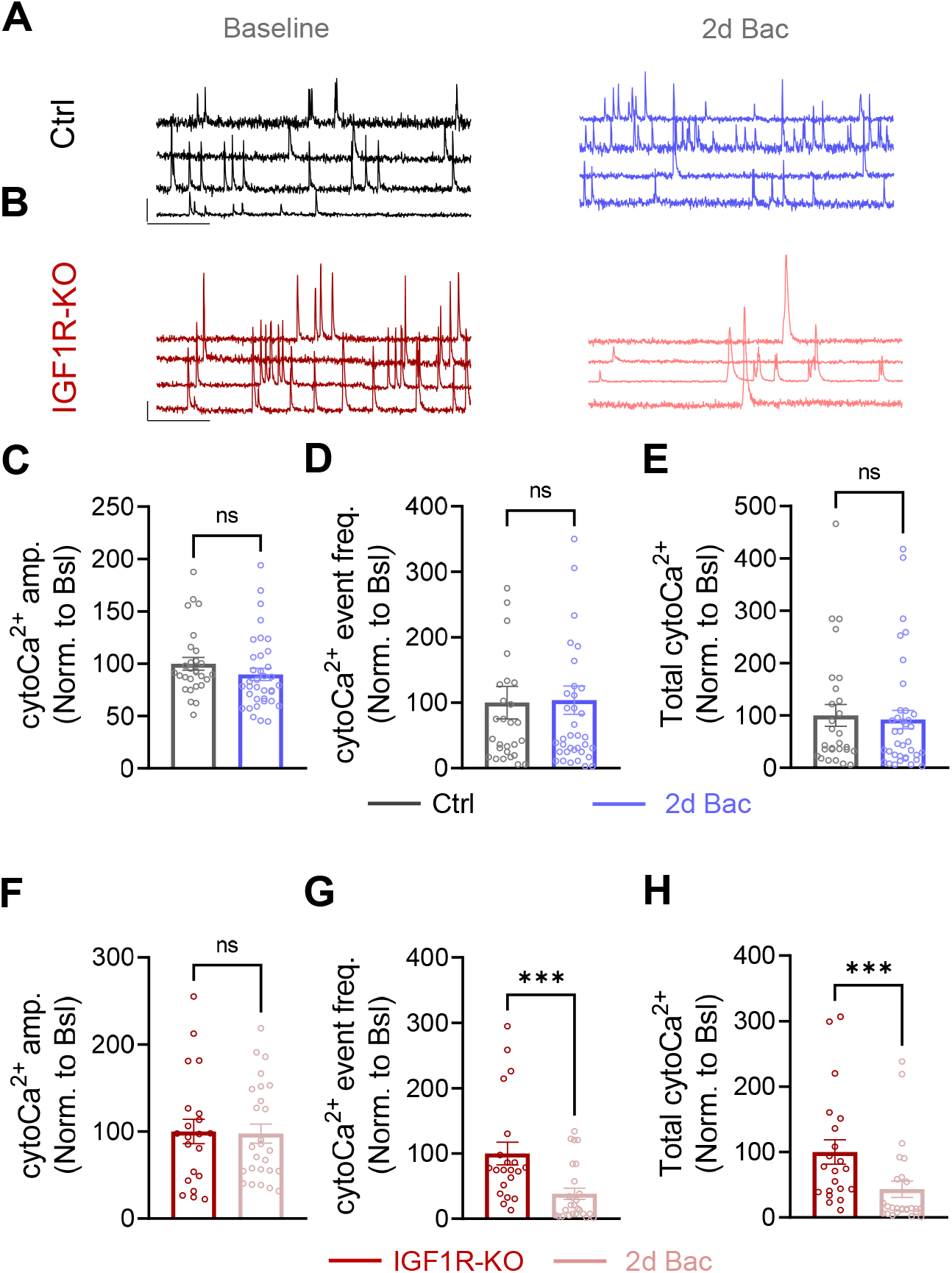
Deletion of IGF1R impairs homeostasis of cytoCa^2+^. (*A*) Example traces of cytoCa^2+^ of Ctrl excitatory neurons at baseline (*Left*) and following two days of Bac (*Right*). Scale bars: 20 sec, 0.5 Δ*F/F*. (*B*) Example traces of cytoCa^2+^ of IGF1R-KO excitatory neurons at baseline (*Left*) and following Bac_2d_ (*Right*). Scale bars: 20sec, 0.5 Δ*F/F*. (*C-E*) Bac_2d_ did not affect mean cytoCa^2+^ amplitude (*P* = 0.07), frequency (*P* = 0.92) and total cytoCa^2+^ (mean amplitude*frequency, *P* = 0.57) in Ctrl neurons (n = 28 for Ctrl; n = 37 for Ctrl+Bac). (*F-H*) Effect of Bac_2d_ in IGF1R-KO neurons (n = 21 for IGF1R-KO, n = 26 for KO+Bac). (*F*) Mean cytoCa^2+^ = 0.92). (*G*) Mean cytoCa^2+^ frequency in IGF1R-KO neurons was reduced to 38.3 ± 8.6% following Bac_2d_ (*P* = 0.0005). (*H*) Total cytoCa^2+^ in IGF1R-KO was reduced to 43.1 ± 12.3% following Bac_2d_ (*P* = 0.0005). Mann-Whitney test. Error bars indicate SEM. ns, not significant, \*\*\**P* < 0.001.

**Figure S4.**
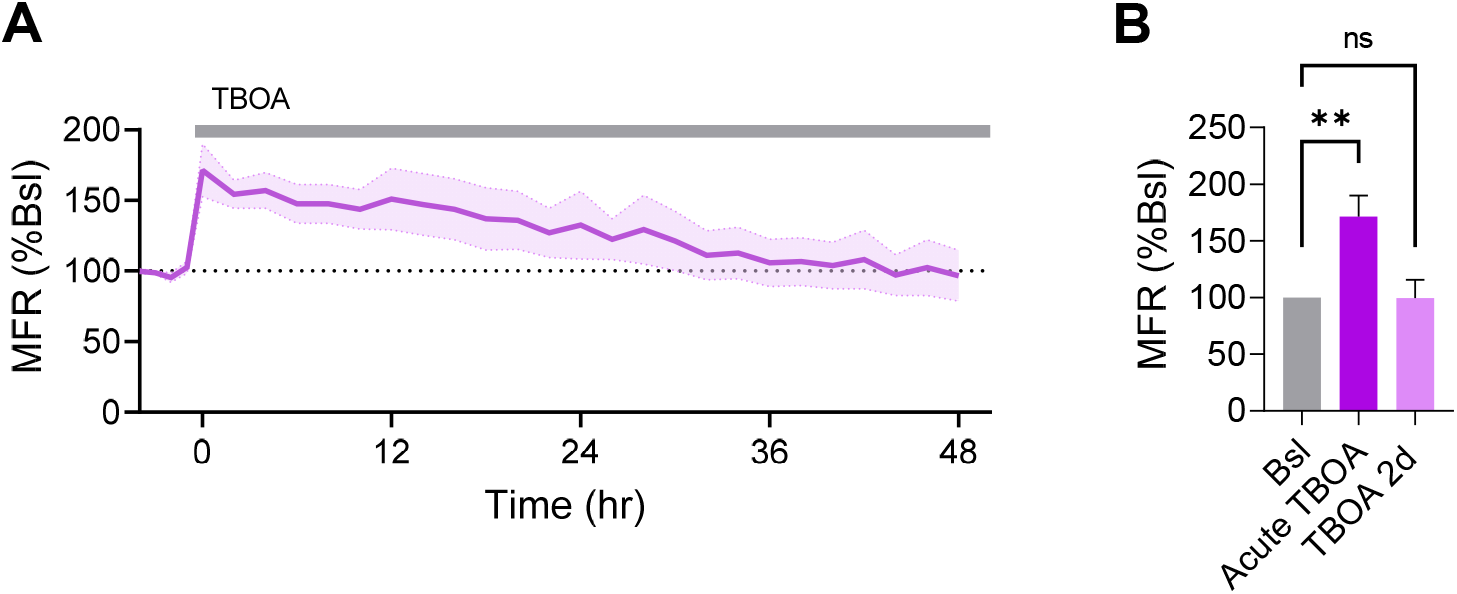
IGF1R-KOs are capable of maintaining MFR homeostasis when activity is increased pharmacologically. *(A)* Mean and SEM of IGF1R-KO cultures, normalized to their baseline firing rate (6 experiments, 430 channels). (*B*) TBOA increased MFR to 171.3 ± 18.6%, followed by a homeostatic re-normalization back to baseline level (*P* > 0.9999, 99.48 ± 16.07 %, N = 6). Friedman’s test with Dunn’s correction for multiple comparisons. Error bars indicate SEM. ns, not significant, \*\**P* < 0.01.

**Figure S5.**
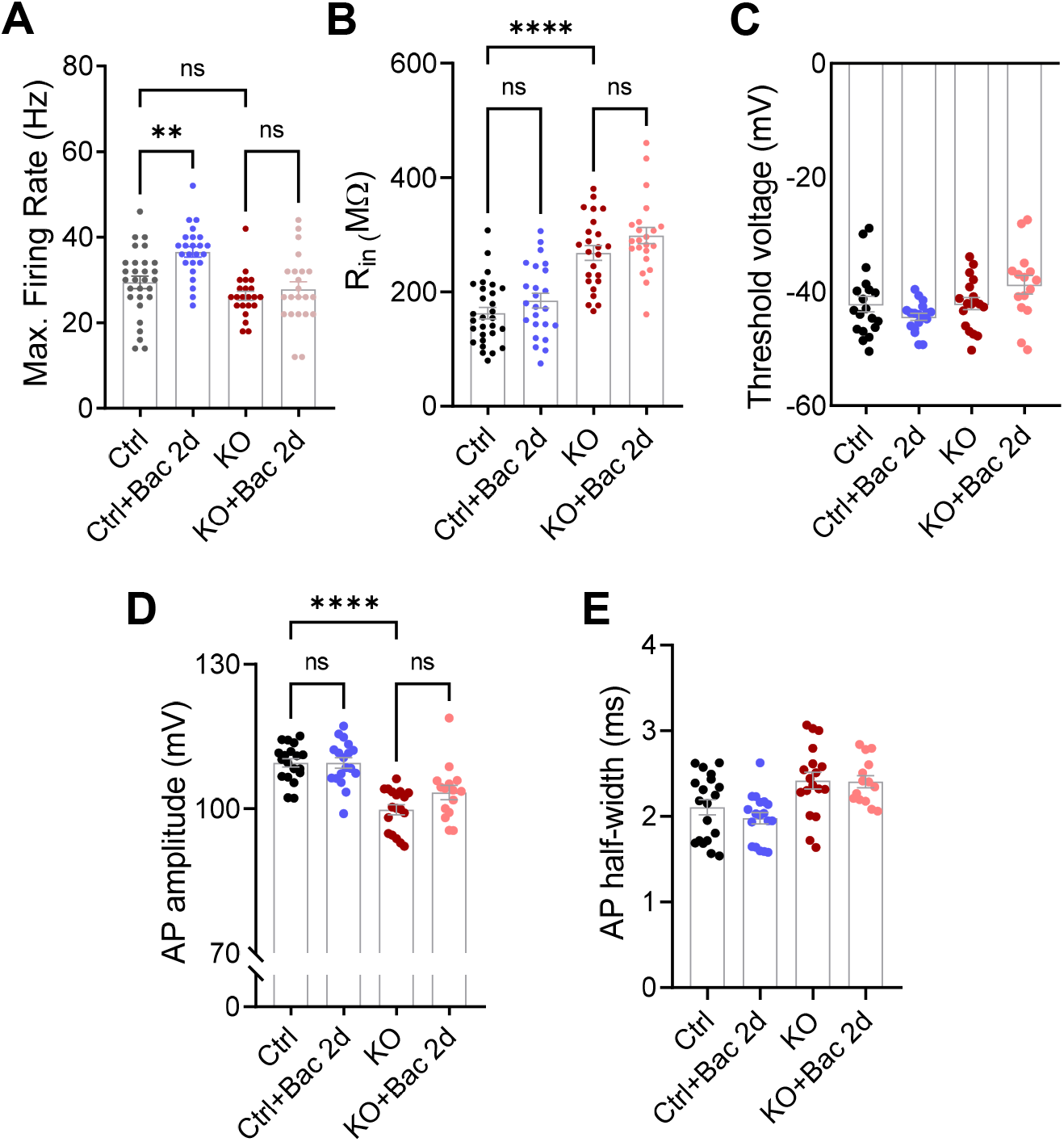
No difference in action potential threshold or waveform, input resistance and mEPSC frequency in either Ctrl or IGF1R-KO following baclofen. (*A*) Maximal firing rate is increased in Ctrl neurons following Bac_2d_ (29.4 ± 1.4 Hz, n = 29 for Ctrl; 36.6 ± 1.2 Hz, n = 24 for Ctrl+Bac), but does not change (*P* > 0.9999) in IGF1R-KO neurons (31.4 ± 2.6 Hz, n = 26 for IGF1R-KO; 27.8 ± 1.8 Hz, n = 22; for KO+Bac). (*B*) No change in input-resistance in neither Ctrl (*P* = 0.864) nor IGF1R-KO (*P* > 0.9999) neurons following Bac_2d_. Increased input resistance in IGF1R-KO neurons compared to Ctrl (162.0 ± 10.3 MΩ, n = 29 for Ctrl; 184.8 ± 12.9 MΩ, n = 24 for Ctrl+Bac; 298.9 ± 24.4 MΩ, n = 26 for IGF1R-KO; 313.8 ± 20.6 MΩ, n = 23 for KO+Bac) (*C*) Threshold voltage does not change for any of the groups (42.19 ± 1.34 mV, n = 19 for Ctrl; −44.42 ± 0.61 mV, n = 18 for Ctrl+Bac; −42.11 ± 1.06 mV, n = 18 for IGF1R-KO; −38.83 ± 1.625 mV, n = 15 for KO+Bac). (*D*) No change in action-potential amplitude in neither Ctrl (*P* > 0.9999) nor IGF1R-KO (*P* = 0.3835) neurons following Bac_2d_. Decreased amplitude in IGF1R-KO neurons compared to Ctrl (109.5 ± 0.87 mV, n = 19 for Ctrl; 109.5 ± 1.083 mV, n = 18 for Ctrl+Bac; 99.77 ± 1.04 mV, n = 18 for IGF1R-KO; 103.4 ± 1.5 mV, n = 15 for KO+Bac). (*E*) No change in action-potential half-width between the groups (2.107 ± 0.087 ms, n = 19 for Ctrl; 1.978 ± 0.063 ms, n = 18 for Ctrl+Bac; 2.418±0.097 ms, n = 18 for IGF1R-KO; 2.406 ± 0.07 ms, n = 15 for KO+Bac). Kruskal-Wallis test with Dunn’s correction for multiple comparisons (*A-E*). Error bars indicate SEM. ns, not significant, \*\**P* < 0.01, \*\*\*\**P* < 0.0001.

**Figure S6.**
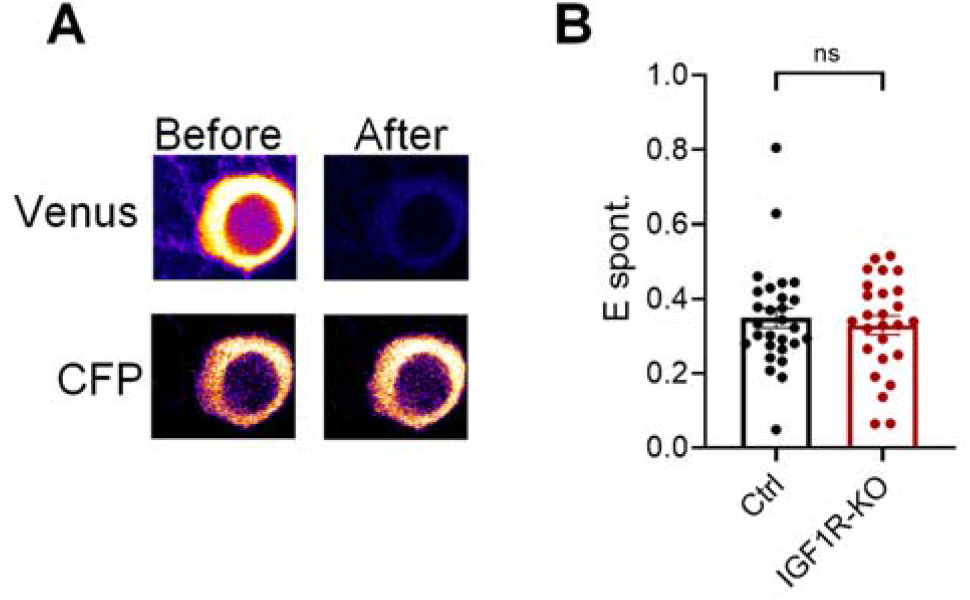
ATP level is unaltered in IGF1R-KO neurons. (A) Representative neuron expressing cytosolic ATeam before (*left*) and after (*right*) acceptor photo-bleaching (*Top*). (B) (C) No difference (*P* = 0.925) in FRET efficiency of spontaneously active Ctrl (0.349 ± 0.026, n = 28) and IGF1R-KO (0.329 ± 0.025, n = 26) neurons. Mann-Whitney test. Error bars indicate SEM. ns, not significant.

**Figure S7.**
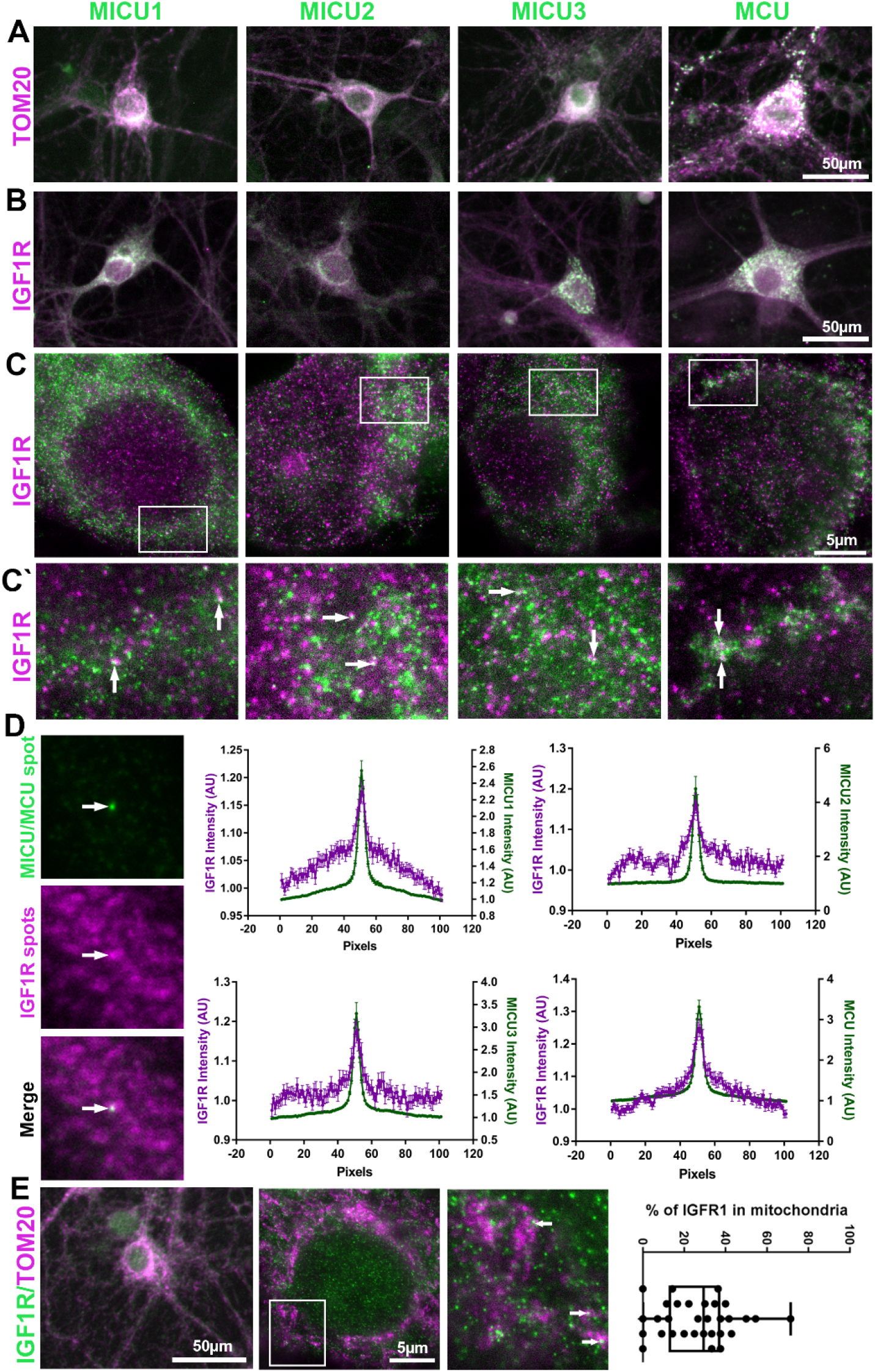
Partial colocalization of the mitochondrial Ca^2+^ uniporter complex (MCUc) and IGFR1 in neurons. (*A*) Hippocampal cultured neurons were immunostained for TOM20, as a mitochondrial marker (*magenta*), in combination with antibodies against four different subunits of the MCUc (*green*). From left to right: MICU1, MICU2, MICU3, and MCU. Scale bars: 50 µm. The images were taken with an epifluorescence microscope. (*B*) Cultured neurons were immunostained as above, to test the colocalization of IGF1R (*magenta*) and the respective MCUC subunits (green). Scale bars: 50 µm. (*C*) Representative STED images of neural cell bodies immunostained for IGF1R (*magenta*) and MCUc subunits (*green*). (*C*’) Enlarged views of the delineated areas alongside the filled arrows depict examples of co-labelling of IGF1R and the MCUc proteins. Scale bars: 5 µm. (*D*) Square regions of interest (ROIs) were obtained for all MCUc spots, both in the MCUc and in the IGF1R channels. The ROIs were then overlaid, which provides a visual indication of the presence of IGF1R in relation to MCUc spots. The arrows point to the ROI centers, where the MCUc spots are located, and where an enrichment of IGF1R is also observed. To analyze these images, line scan profiles of intensity for IGF1R (*magenta*) and the four MCUc subunits (*green*) were performed across the ROIs, and are shown as means ± SEM from 8 samples for each MCUc subunit. A small, but significant enrichment of IGF1R near MCUc spots can be observed for all samples (Kruskal-Wallis tests followed by post-hoc Tukey test, corrected for multiple comparisons; *P* < 0.001 for colocalization with MCU, MICU1 and MICU2; *P* < 0.05 for MICU3). (E) We immunostained the cultured neurons for IGF1R (green) and the mitochondrial marker TOM20 (magenta), to estimate the proportion of IGF1R that colocalizes with mitochondria. STED images strongly suggest a partial colocalization in the soma (some colocalized molecules are shown by the arrows). Scale bars, 50 µm and 5 µm, respectively. The quantification shows the percentage of IGF1R spots found within the space delimited by the TOM20 staining (i.e., found on mitochondria), which averages to ~30%. The box shows the median and the quartiles, while the whiskers show the range of data (N = 5 independent experiments, with 5-7 cells imaged for each experiment).

## Supplementary Methods

### STED microscopy

Immunostaining was carried out based on standard protocols. Cultured hippocampal neurons (prepared exactly as described (75)) were fixed with 4% PFA in PBS for 30 min, followed by quenching with 100 mM glycine in PBS for 15 min, before being permeabilized and blocked using a blocking buffer prepared with 2% BSA (Sigma, 9048-46-8) in PBS added to 0.1% TritonX-100 (Merck, Kenilworth, NJ, USA). Next, the cells were incubated overnight with primary antibodies diluted in blocking reagent, at 4 °C. The neurons were then incubated with the appropriate secondary antibody (1:400) for 60 min at room temperature. The coverslips were mounted using Mowiol (Merck Millipore, Kenilworth, NJ, USA). The following primary antibodies were used: MICU1 rabbit polyclonal (1:100; Sigma), MICU2 rabbit polyclonal (1:50; Sigma), MICU3 rabbit polyclonal (1:150; Sigma), MCU rabbit polyclonal (1:100; Sigma), TOM20 mouse monoclonal (1:100; Sigma), and IGF1R mouse monoclonal (1:100; Thermofisher). The applied secondary antibodies were anti-mouse STAR580 and anti-rabbit STAR635P purchased from Abberior GmbH, Göttingen, Germany.

Epifluorescence images (Fig. 4A, B) were obtained by means of an IX83 inverted microscope (Olympus). STED imaging (Fig. 4C,C’) was captured using a STED Abberior microscope, Göttingen, Germany. Excitation lines of 640 nm and 561 nm were adopted for exciting Star653P (MCUc subunits) and Star580 (TOM20 and IGF1R). For STED excitation, pulsed lasers were set at 640 nm and 580 nm. STED depletion was implemented via 775 nm depletion laser and the images were acquired at 20 nm pixel size.

### Western Blot

Primary hippocampal cultures were infected on DIV5-6 by AAVs encoding cre-P2a-mCherry sequences (+Cre) or P2a-mCherry sequence (-Cre). Lysates of equal number of cells (~500k) were separated by SDS-PAGE, transferred to nitrocellulose, blocked with 5% skim milk powder (Difco # 232100) and incubated ON with anti-IGF-1R antibody (#3027 CST), followed by anti-rabbit-HRP secondary antibody (#11-035-044 Jackson ImmunoResearch). To normalize the signal, the blot was re-probed with anti-actin antibody (Sigma), followed by anti-mouse-HRP secondary antibody (#115-035-146 Jackson ImmunoResearch).

### Knockout validation at the level of genomic DNA

Primary hippocampal cultures were infected on DIV5-6 by AAVs encoding cre-P2a-mCherry sequences (+Cre) or P2a-mCherry sequence (-Cre). Cells were collected and lysed, and DNA was purified using the MasterPure™ DNA&RNA Complete purification kit according to the company’s manual (Cat. MC85200 epicentre). DNA was probed with primers for the excised and not excised version of the floxed exon3 of the IGF1R gene with the following primers:

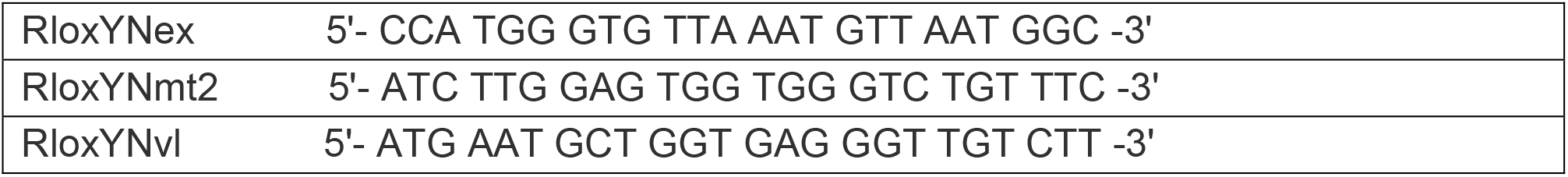

### FRET efficiency imaging

Primary hippocampal cultures were infected on DIV5-6 by AAV1/2 encoding Cre-P2a-mCherry sequences or P2a-mCherry sequence. Experiments were performed on a FV-1000 (Olympus, Japan) system at room temperature. Some coverslips were placed in Tyrode’s solution that contained (in mM): NaCl, 145; KCl, 3; glucose, 15; HEPES, 10; MgCl_2_, 1.2; CaCl_2_, 1.2; pH adjusted to 7.4 with NaOH. Others were placed in their own media, in a heated (34-35°C) Stage-Top Incubator System TC, connected to a CU-501 temperature controller and a humidifier delivering 5% CO_2_ air (Live Cell Instruments, Republic of Korea). The results were similar for both conditions so they were pooled. Infection of neurons with AAV1/2-CBAP-Cre-mCherry or AAV1/2-CBAP-mCherry was verified with a 561 nm laser. Images of neuronal soma were taken before and after acceptor (cpmVen) photobleaching. Donor (mseCFP) was excited with a 440 nm laser, and emission was measured at [460-500] nm before (I_DA_) and after (I_D_) photobleaching. Bleaching was accomplished using a 514 nm laser. Images of acceptor were taken before and after bleaching at [530-600] nm, to asses bleaching level: neurons with less than 85% reduction in fluorescence were excluded. FRET efficiency was calculated as [I_DA/_ I_D_].

## Notes

### Competing Interest Statement

The authors have declared no competing interest.

